# Lignature: A Comprehensive Database of Ligand Signatures to Predict Cell-Cell Communication

**DOI:** 10.1101/2025.03.22.644770

**Authors:** Ying Xin, Md Amanullah, Cheng Qian, Chingmo Zhou, Jiang Qian

## Abstract

Ligand-receptor interactions mediate intercellular communication, inducing transcriptional changes that regulate physiological and pathological processes. Ligand-induced transcriptomic signatures can be used to predict active ligands; however, the absence of a comprehensive set of ligand-response signatures has limited their practical application in predicting ligand-receptor interactions. To bridge this gap, we developed Lignature, a curated database encompassing intracellular transcriptomic signatures for 362 human ligands, significantly expanding the repertoire of ligands with available intracellular response signatures. Lignature compiles signatures from published transcriptomic datasets and established resources such as CytoSig and ImmuneDictionary, generating both gene- and pathway-based signatures for each ligand. We applied Lignature to predict active ligands driving transcriptomic changes in controlled *in vitro* experiments and real-world single-cell sequencing datasets. Lignature outperformed existing methods such as NicheNet, achieving higher accuracy in identifying active ligands at both the gene and pathway levels. These results establish Lignature as a robust platform for ligand signaling inference, providing a powerful tool to explore ligand-receptor interactions across diverse experimental and physiological contexts.

## INTRODUCTION

Cell-cell communication is essential for coordinating biological processes in multicellular organisms, with ligand-receptor (LR) interactions serving as a primary mechanism in this process^1–3^. The rapid advancement of sequencing technologies has enabled the development of numerous computational methods for predicting potential LR interactions from transcriptomic data.

Current methods for predicting cell-cell communication, particularly LR interactions, fall into three major categories. The first category identifies LR interactions based on the co-expression of ligand genes in sender cells and receptor genes in receiver cells. Tools such as CellPhoneDB^4^, CellChat^5^, SingleCellSignalR^6^, NATMI^7^, and ICELLNET^8^ exemplify this approach. While these methods are effective, they often generate a high number of false positives by relying solely on ligand and receptor gene expression levels.

To enhance accuracy, a second type of approach integrates intracellular networks to model the impact of ligands on target genes in receiver cells. Tools like CCCExplorer^9^, NicheNet^10^, SoptSC^11^, and LRLoop^12,13^ predict LR interactions by comparing observed transcriptional changes with predicted target genes associated with each ligand. However, the incomplete and potentially noisy nature of current signaling and regulatory networks leads to inaccuracies in target gene prediction. Additionally, these methods fail to model the direction and magnitude of target gene expression changes in response to specific ligands.

A third group of methods, including SpaOTsc^14^ and COMMOT^15^, incorporates spatial information to better account for potential signaling between cells, further refining predictions of cell-cell communication. However, these methods rely on spatial transcriptomics data and do not resolve the inherent challenges of expression-based prediction.

In this work, we propose a new approach to predict the specific ligands responsible for the transcriptomic changes within the receiver cells. Instead of merely modeling the target genes of ligands, our method will identify the signatures of ligands by measuring the transcriptomic changes derived from actual gene expression data. When cells are treated with a given ligand, and their gene expression is measured before and after treatment, we can then create a distinct signature unique to that particular ligand. It’s worth noting that the concept of ligand signatures is widely accepted in the field. Many researchers have identified the “responsive genes” of the ligands^16–20^. Previous works such as CytoSig^21^ and ImmuneDictionary^22^ indeed collect the signatures of ligands. However, they focus exclusively on cytokines, limiting the practical applications of their datasets to the prediction of cell-cell communication. Here we provide a comprehensive ligand signature database, Lignature, which contains the signatures for 362 ligands across diverse categories. Additionally, we constructed gene- and pathway-based signatures for these ligands. We validated these signatures using in vitro experiments and single-cell sequencing datasets, demonstrating their reliability. The accompanying computational tool further enables the signature-based ligand-receptor interaction predictions.

## RESULTS

### Overall design

The transcriptomic changes induced by a specific ligand are referred to as the ligand’s signatures. By comparing observed gene expression changes to a comprehensive library of ligand signatures, it is possible to predict which ligands are acting on the cells, or which ligands are likely to induce observed gene expression changes. To achieve this, we collected transcriptomic datasets where gene expression profiles were measured before and after treatment with specific ligands. These datasets were used to construct a library of ligand signatures based on the resulting transcriptomic changes.

Recognizing that ligand-induced signatures can vary depending on cell types and experimental conditions, we included multiple signatures for a given ligand when such datasets were available. Additionally, we developed a companion software tool designed to predict and visualize active ligands by leveraging this curated collection of ligand signatures.

### Generation of the Lignature database

We developed an automatic pipeline to search Gene Expression Omnibus (GEO)^23^ for datasets with an experimental design that would potentially contain transcriptomic responsive to these ligands. Starting with the metadata of all 25,868 platform identifiers downloaded from GEO, we collected metadata for 72,738 data-series corresponding to Homo sapiens platforms and molecule type “total RNA”, “polyA RNA”, or “nuclear RNA”, where each data-series could include multiple datasets. The curated metadata of the GEO data-series was then queried with a predefined set of 859 ligands based on our previous work LRLoop^12^. We searched the metadata with ligand genes’ full names, symbols, and aliases, together with a set of 105 experiment keywords such as “ligand”, “receptor”, “treatment”, “stimulate”, “overexpress”, “block”, “knockdown”, and “silence”, resulted in a list of 9,187 data-series for manual curation. Finally, after carefully examining the description for each of the collected data-series in GEO, we removed those that did not serve our purpose, resulting in a list of 732 data-series, containing transcriptomic response data for 281 ligands that were not included in the database CytoSig^21^ (Figure 1a).

**Fig.1.**
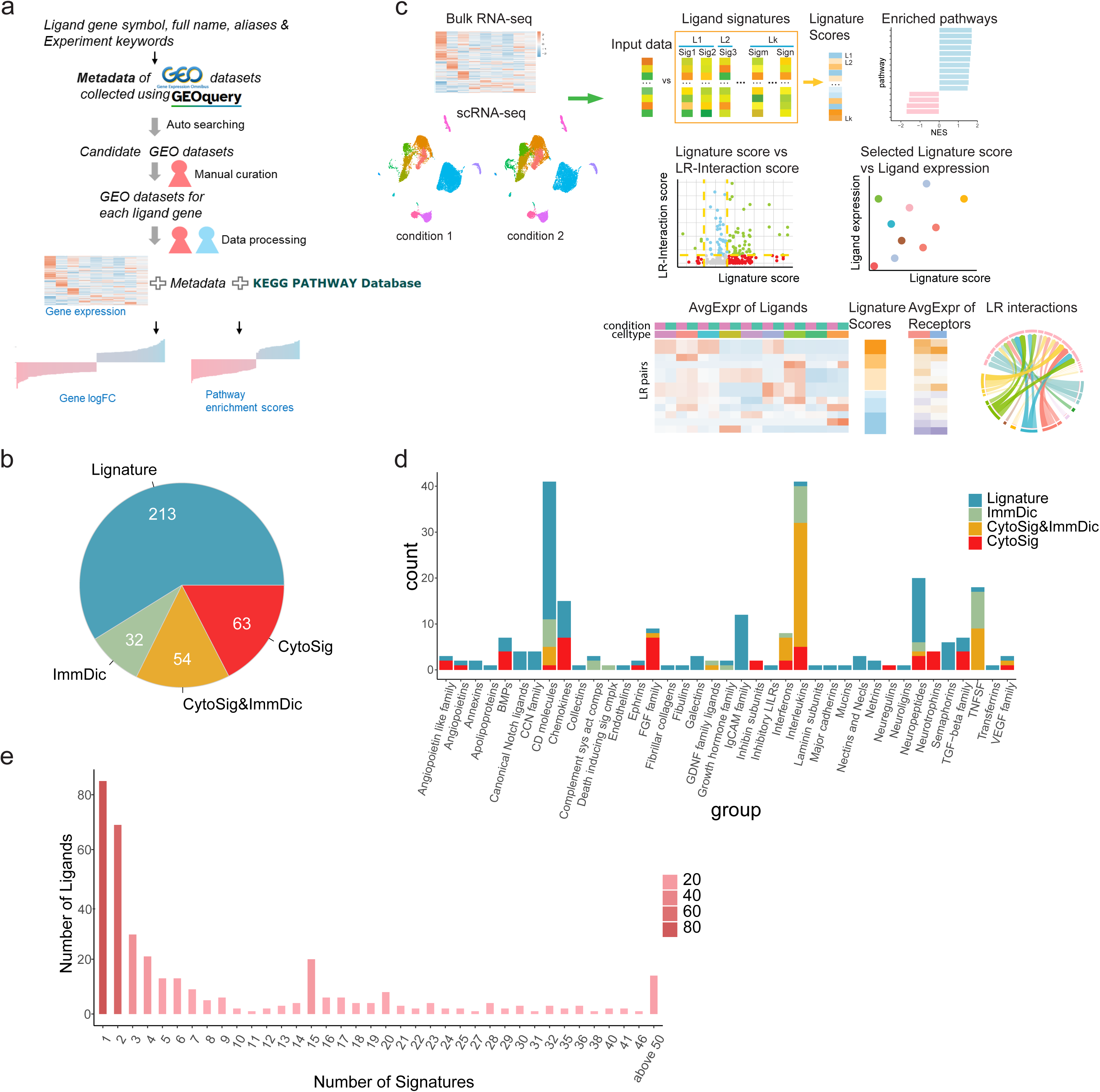
Curation and overview of Lignature. **(a)** Flowchart of generating the Lignature database. **(b)** Number of the ligands collected in the Lignature database from different data sources. **(c)** Schematic diagram of application of Lignature in predicting and visualizing active LR interactions based on ligand signatures and LR expression. **(d)** HUGO Gene Nomenclature Committee categories of the ligands curated in the Lignature database. **(e)** Distribution of the number of signatures of each ligand in Lignature.

The transcriptomic datasets, including gene expression array and RNA-seq datasets, were then processed for differential expression analysis (see Methods for details), vectors of log2FC (fold change) values for expressed genes were collected as the gene-level ligand signatures. For datasets with multiple treatment conditions (e.g., durations or doses), a separate signature was created for each different condition. After combining the data of 149 ligands from CytoSig^21^ and ImmuneDictionary^22^, we obtain the datasets for 362 ligands (Figure 1b). In addition to the gene-level response signatures, we also calculated KEGG pathway signatures for each ligand. Specifically, for each gene-level signature, we performed Gene Set Enrichment Analysis for 302 KEGG pathways with 10 ∼ 500 genes, calculated the Normalized Enrichment Score (NES) for each KEGG pathway. The vectors of NES were collected as the pathway-level signatures (Figure 1a).

Using the Lignature database as a reference, we provided a companion computational tool that infers the ligands responsible for transcriptomic changes within receiver cells, and the corresponding cell-cell interaction networks between multiple cell types or clusters. For a dataset of interest, Lignature scores each ligand in the database by the similarity of their corresponding signatures compared to the gene expression profile from the input data. The tool provides multiple options for calculating expression-based LR interaction strength scores and visualization, providing an intuitive summary of the predicted LR activities across different conditions (Figure 1c).

### Overview of the Lignature database

The 362 ligands are classified into 40 groups based on the HUGO Gene Nomenclature Committee categories^24^. The largest ligand groups include CD molecules, interleukins, neuropeptides, transforming growth factor beta (TGF-beta) superfamily, and chemokine ligands (Figure 1d). This classification also highlights data source biases; CytoSig and ImmuneDictionary primarily focus on interleukins, TNFSF, interferons, and FGF family, while our newly curated data cover a broader range, including most CD molecules, neuropeptides, Ig-like cell adhesion molecules (IgCAM), and semaphorins. Additionally, 153 ligands in Lignature are not part of any of these 40 groups.

A single ligand may have multiple signatures, as it may produce different responses in various cell types with diverse treatment durations and concentrations. In Lignature, the number of signatures per ligand ranges from 1 to 271, with TGFB1 having the largest number of signatures (Figure 1e). We systematically compared the similarity of signatures of the same ligand with those from different ligands by calculating Pearson correlation coefficients for all signature pairs (Figure 2a). The mean correlation coefficients between signatures of the same ligand are 0.174, 0.161, and 0.064 in Lignature, CytoSig, and ImmuneDictionary, respectively. In contrast, the mean correlation coefficients between signatures of different ligands are 0.003, 0.022, and 0.058 for Lignature, CytoSig, and ImmuneDictionary, respectively. Although diverse responses could be induced by the same ligands in different cell types, signatures of the same ligand still exhibit significantly higher similarity than those of different ligands (Figure 2a).

**Fig.2.**
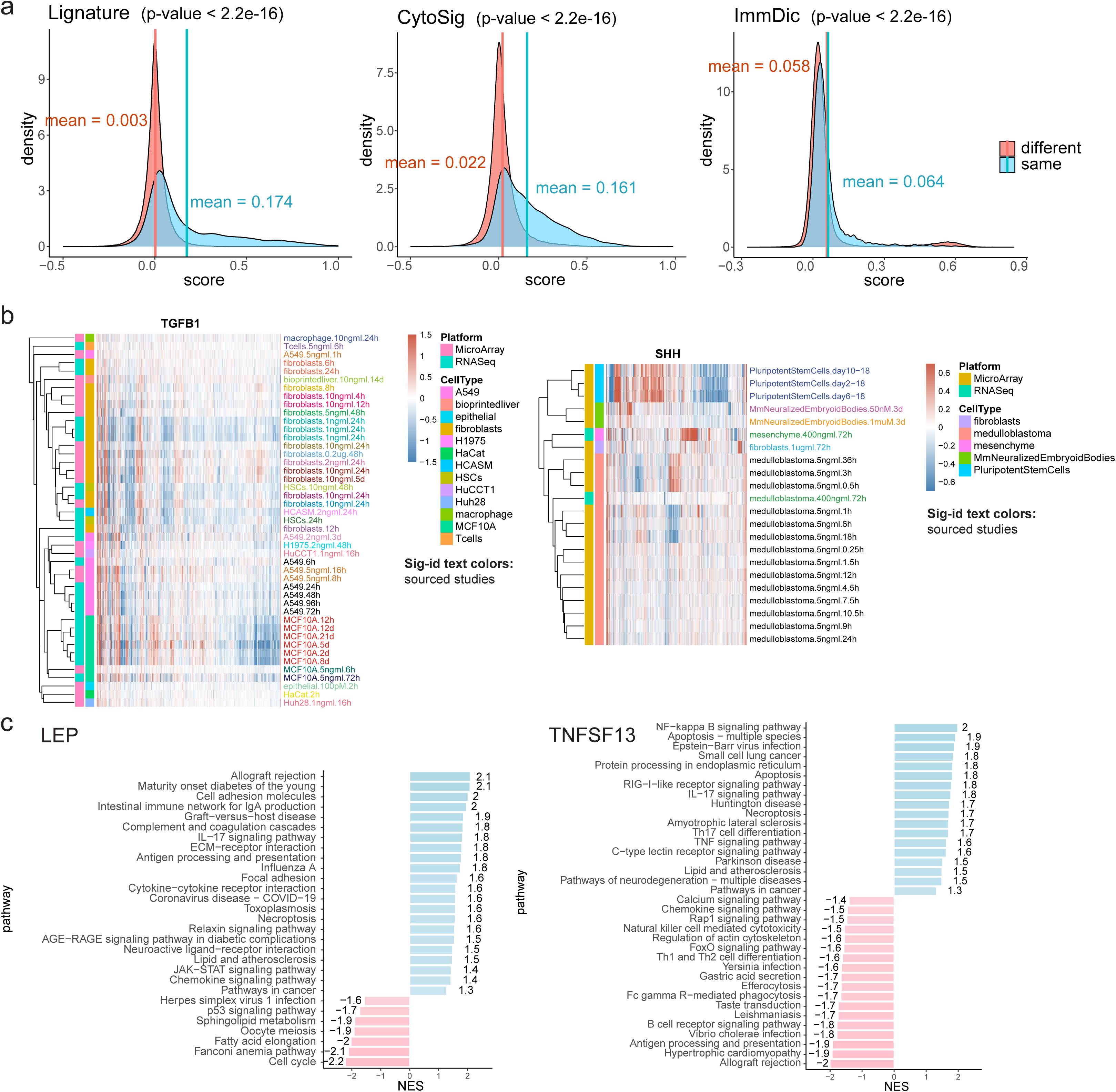
Properties of ligand signatures in Lignature. **(a)** Pearson correlation coefficients of gene-level signatures of the same ligand are significantly higher than signatures from different ligands. P-values were calculated using KS-test. **(b)** Heatmaps of sample gene-level signatures of SHH and TGFB1 exhibit diversity in signatures for individual ligands. **(c)** Examples of enriched up- and down-regulated KEGG pathways in sample ligand signatures (LEP and TNFSF13).

We further analyzed the diversity of signatures for individual ligands. For example, TGFB1 signatures show that the signatures of the same cell types or cell lines (e.g., fibroblasts, A549, MCF10), from the same or different studies, tend to cluster together, with overall patterns remaining conserved regardless of the data platform (Figure 2b). Similarly, SHH has over twenty signatures from datasets across different organisms, cell types, treatment methods, doses, durations, and platforms. The signatures from the same cell types, such as mouse neuralized embryoid bodies and medulloblastoma, also cluster together, including those from different studies (Figure 2b). This analysis suggests that the cell type is one major source of diversity in signatures for the same ligands.

To confirm the biological relevance of the ligand signature, we calculated the enrichment of up- and down-regulated genes of ligand signatures in KEGG pathways. For instance, the enriched pathways for the signature genes of ligand LEP (which encodes leptin) include allograft rejection, intestinal immune network for IgA production, graft-versus-host disease, complement and coagulation cascades. These pathways are related to LEP’s involvement in inflammatory processes^25–27^. Conversely, the p53 pathway is enriched in down-regulated genes, consistent with leptin’s inhibitory effect on cell cycle processes by decreasing the key p53 proliferation regulator^28^ (Figure 2c). Similarly, the signature genes of ligand TNFSF13 shows enrichment of upregulated genes in NFkB signaling, apoptosis, TNF signaling pathway and other pathways that are highly consistent with the known cellular functions of TNFSF13^29^. It is worth noting that TNFSF13 is also closely associated with allograft rejection in organ transplantation^30^ (Figure 2c, for more examples, see SupplFigure 1). These enriched pathways can be used to constructed pathway-based signatures and predict the active ligands.

### Assessment in signaling inference with bulk *in vitro* datasets

We assessed the ability of Lignature to predict active ligands using ligand signatures. To this end, we collected and processed additional transcriptomic datasets as test cases, where individual ligands were used to treat cell lines, and transcriptomic changes were measured before and after treatment. The goal was to predict the acting ligands based on these observed transcriptomic changes. A total of 90 test datasets were used, including 24 human and 66 mouse datasets, representing 27 unique treatment ligands (Figure 3a). To ensure comparability, genes from mouse datasets were converted to human gene symbols using their human homologs.

**Fig.3.**
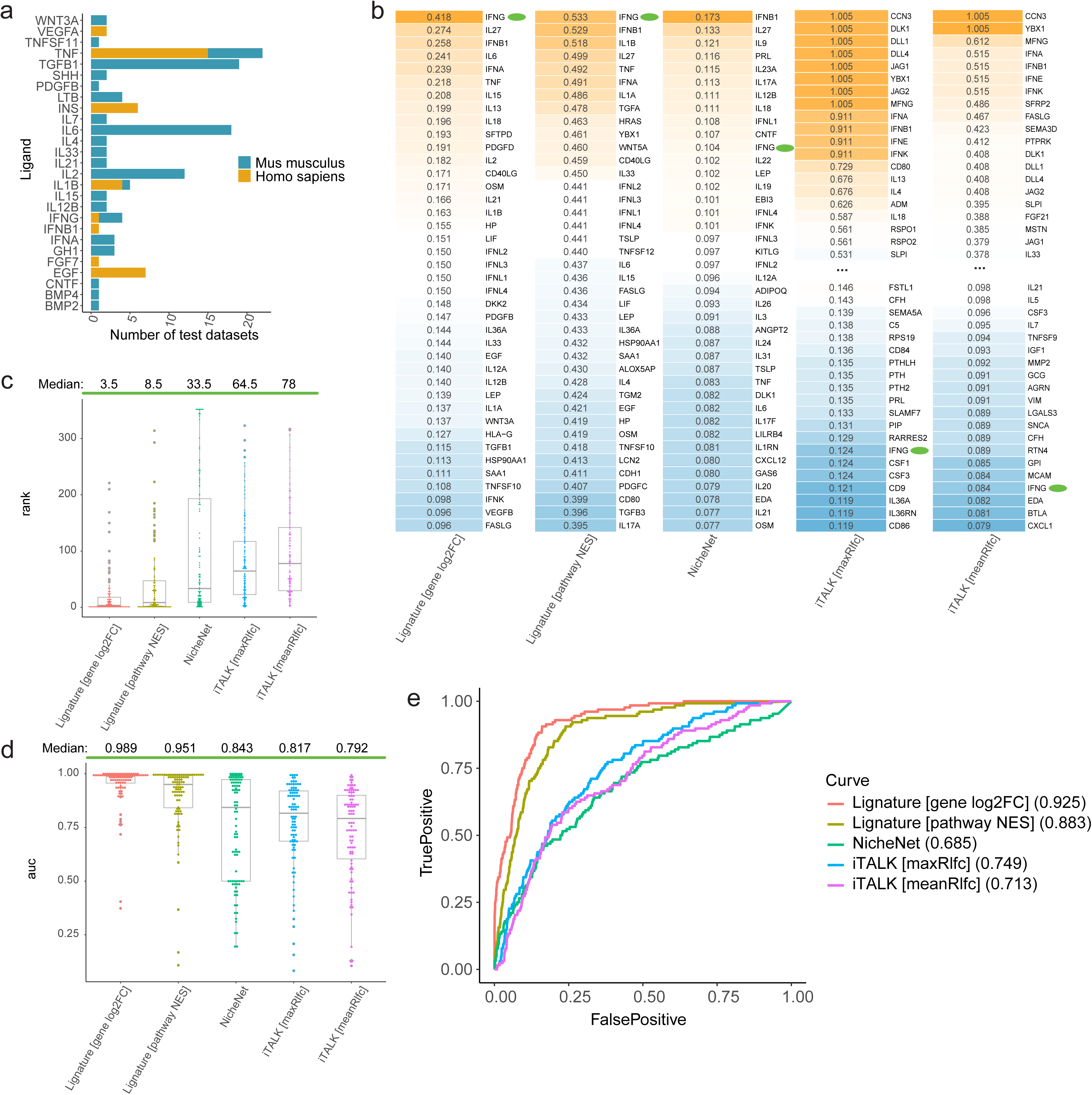
Comparison of Lignature to other methods in signaling inference with bulk in vitro datasets. **(a)** A total of 90 transcriptomic datasets of ligand-treated cell lines used as test datasets, including 24 human and 66 mouse datasets representing 27 unique treatment ligands. **(b)** Top ranked ligands (among all ligands included in Lignature) and the corresponding ligand scores of an example test dataset treated with IFNG, predicted from gene-level Lignature, pathway-level Lignature, and three other methods. **(c)** Ranks of the true treatment ligands among all ligands included in Lignature of all 90 test datasets predicted by each method. **(d)** AUC values of the ROC curves for predictions of the 90 test datasets from each method. **(e)** ROC curves and the corresponding AUC values for combined predictions of all the 90 test datasets from each method.

For each test dataset, we calculated a similarity score between the observed transcriptomic changes and signatures in the Lignature library. The signatures were ranked based on these similarity scores, and the ranked position of the treatment ligand was examined. For example, in a dataset treated with ligand IFNG, the transcriptomic changes were compared against the signatures of the ligands in Lignature. Both the gene-level and pathway-level signatures for ligand IFNG exhibit high correlation coefficients with the test data, accurately identifying it as an acting ligand responsible for the observed transcriptomic changes (ranked number 1; Figure 3b). Three other methods for ligand activity prediction were compared to Lignature. For the same test dataset, NicheNet ranked IFNG number 11, lower than the prediction from Lignature yet a reasonably high rank for identifying IFNG as a treatment ligand. Another method iTALK that predicts LR interactions based on differential expression of receptor genes alone ranked IFNG much lower at number 279 and 306, respectively, when scoring each ligand by the maximum or mean absolute log2FC values of its cognate receptors (denoted by maxRlfc and meanRlfc) (Figure 3b).

Across all 90 test datasets, the median rank of treatment ligands was 3.5 and 8.5 for predictions from Lignature gene- and pathway-level signatures (Figure 3c), the median AUC values of the corresponding ROC (receiver operating characteristic) curves are 0.989 and 0.951 (Figure 3d), and the AUC values of the combined ROC curves from predictions for all 90 test datasets are 0.925 and 0.883, respectively (Figure 3e). Applying NicheNet, iTALK maxRlfc, and iTALK meanRlfc to the same datasets yielded median ranks of 33.5, 64.5, and 78 for the treatment ligands (Figure 3c), median AUC values of the corresponding ROC curves 0.843, 0.817, and 0.792 (Figure 3d), and AUC values of combined ROC curves 0.685, 0.749, and 0.713 (Figure 3e). In summary, Lignature demonstrates significantly better performance than the other existing methods.

We next asked whether the signatures from one species can be applied to another. Since Lignature contains mostly signatures from human datasets, we compared its performance on 24 human and 66 mouse test datasets to determine whether human datasets yield better results (SupplFigure 2). For the human test datasets, the median rank of treatment ligands was 2 for both Lignature gene- and pathway-level signatures, while the corresponding median rank for the mouse test datasets are 4 and 13 for gene- and pathway-based signatures. Comparable results were obtained for human and mouse test datasets, demonstrating that the collection of signatures in Lignature is mostly evolutionarily conserved and transferable between human and mouse datasets in the inference of active ligands.

### Application and assessment of Lignature with scRNA-seq datasets

While we assessed the overall performance of ligand signatures using the well-controlled *in vitro* dataset in the previous section, here we applied Lignature to two scRNA-seq datasets that preserved more natural *in vivo* conditions involving multiple cell types.

#### Example 1: SHH-Induced Thalamic Differentiation

To investigate SHH-mediated thalamic differentiation, organoids were treated with recombinant SHH, and scRNA-seq was performed on samples collected on day 70 with or without SHH treatment^31^. Major cell clusters were identified, including excitatory neurons (ExNs), inhibitory neurons (INs), astrocytes (ASs), endothelial cells (ECs), and ependymal cells (Epn) (Figure 4a-b). Since SHH activation promotes the generation of INs, we focused on the IN cluster from the SHH day 14–22 treatment sample compared to the untreated sample.

**Fig.4.**
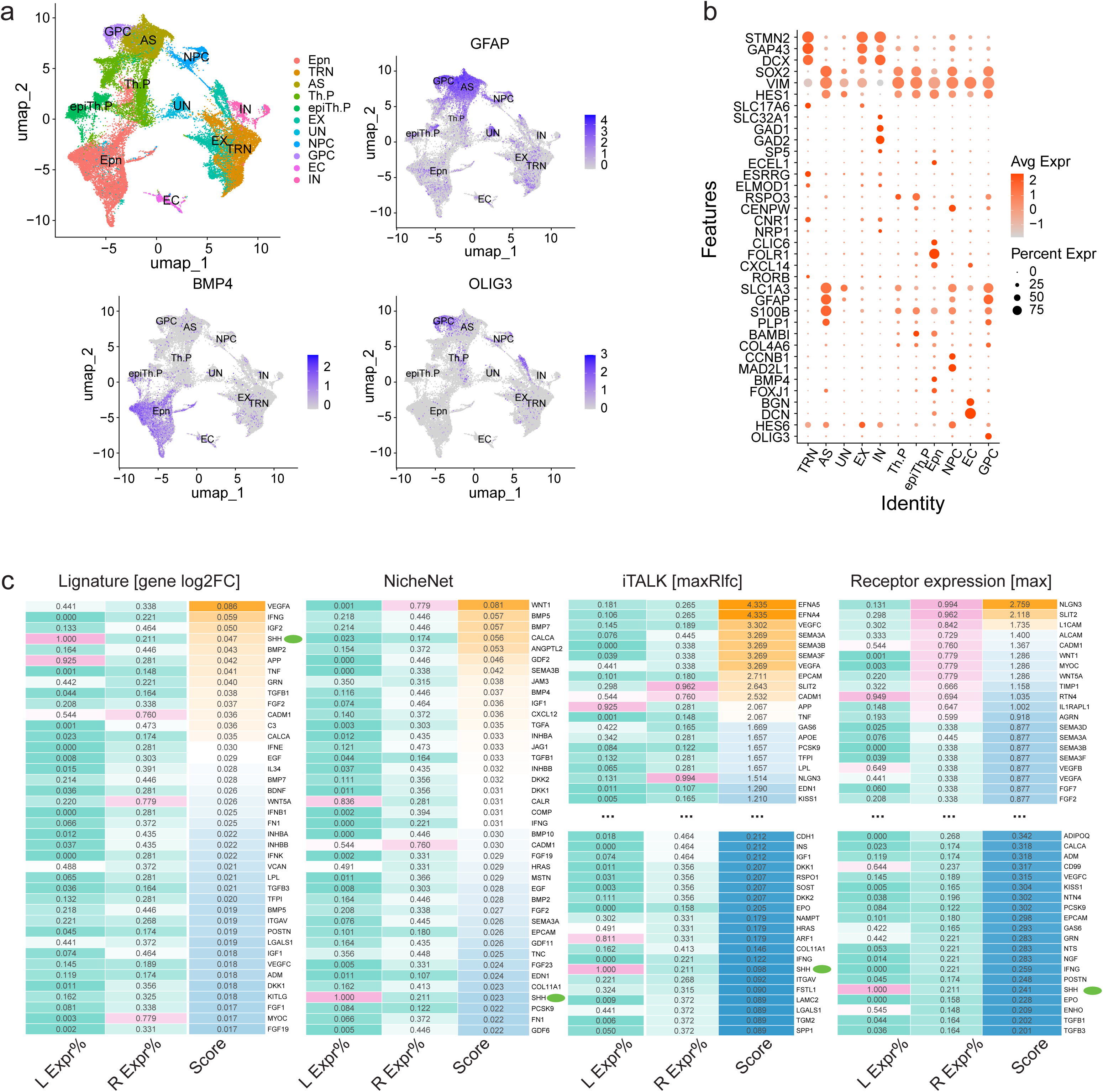
Application and assessment of Lignature on SHH-induced thalamic differentiation scRNA-seq dataset. (a-b) UMAP dimensionality reductions, example feature plots, and DotPlot of selected marker genes of major cell clusters of the SHH-induced thalamic differentiation scRNA-seq dataset. **(c)** Heatmaps of top predicted ligand scores from Lignature and three other methods, together with detection rate of the ligands in all cells and the maximum detection rate of their cognate receptors in IN cells.

Candidate ligands in Lignature were first filtered based on the detection rate of their receptors, retaining ligands with at least one receptor detected at a rate above 0.1. Using Lignature (gene-level signatures), SHH was ranked as one of the top ligands (number four) predicted to be active inducing the observed transcriptomic changes (green dots in Figure 4c). Three ligands, VEGFA, IFNG, and IGF2 ranked higher than SHH. Among them, IFNG can be easily ruled out as false positive, because the expression level of IFNG in all cells are zero (Figure 4c). VEGFA and IGF2 are known to be the downstream regulators of SHH, which play a role in neurogenesis, neuroprotection, and the processes of synaptic formation throughout neural development^32–36^. This result suggests that Lignature captures the transcriptomic response from the signaling cascade initiated by SHH. In contrast, NicheNet, iTALK (maxRlfc) and a receptor expression-based method (maximum expression of the cognate receptors) ranked SHH much lower at number 37, 114 and 126, respectively. In summary, Lignature successfully identified SHH as an active ligand driving transcriptomic changes between the treated and untreated samples, demonstrating its superior accuracy over other methods in identifying ligand activity during induced IN generation.

#### Example 2: Microglia and Müller Glia (MG) Regeneration

Microglia cells have been implicated in suppressing Müller glia (MG) regeneration capacity, as microglia ablation improved retinal neurogenesis from MG^37^. Furthermore, treatment with TNF-α in microglia-ablated mice suppressed the enhanced neurogenesis observed in the absence of microglia^37^, suggesting that TNF-α expressed in microglia might inhibit MG regeneration. scRNA-seq was performed on microglia and MG before and after microglia ablation^37^ (Figure 5a-b), and we aimed to predict the ligands responsible for the transcriptomic changes in MG.

**Fig.5.**
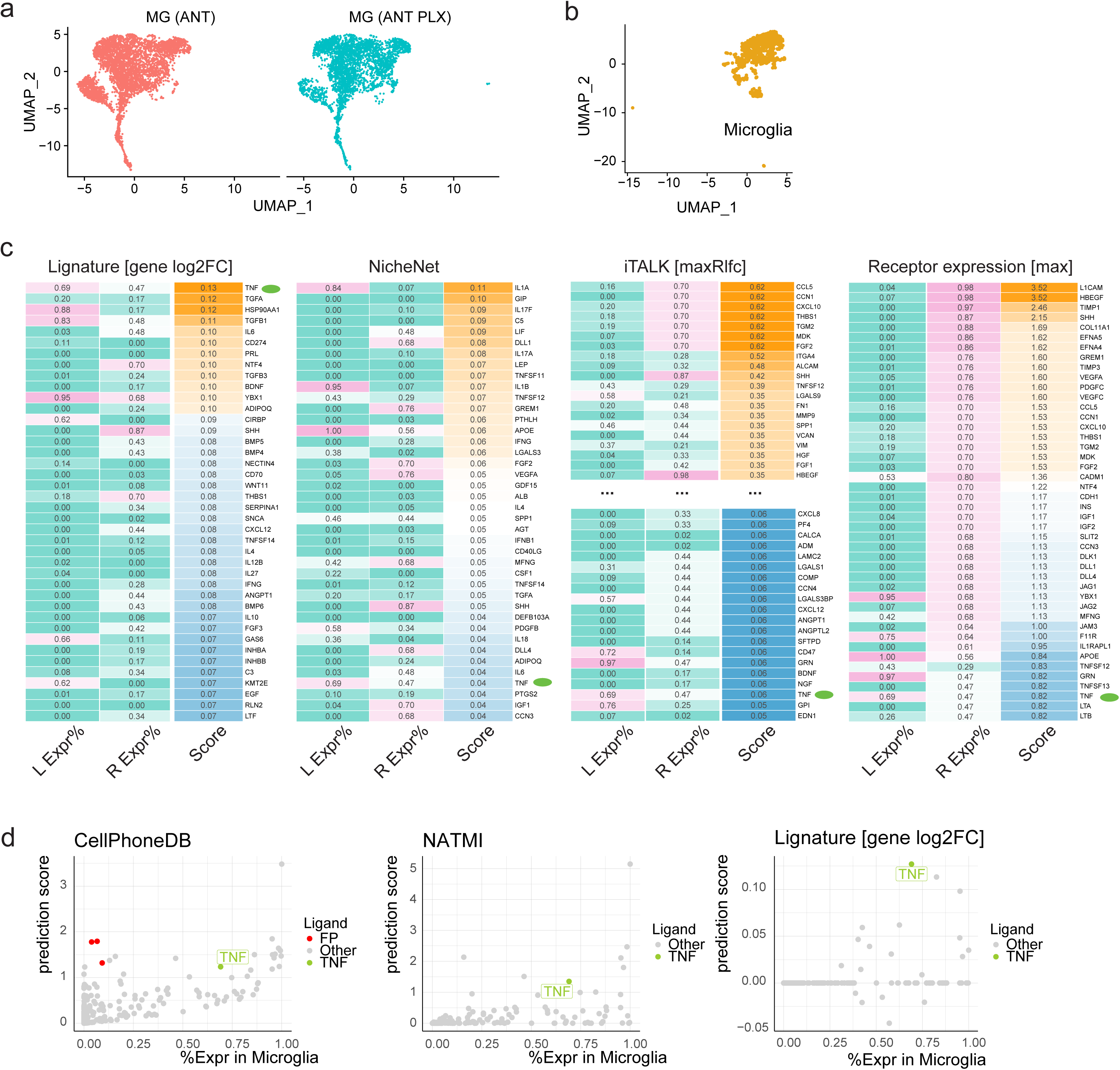
Application and assessment of Lignature on microglia and Müller glia regeneration scRNA-seq dataset. (a-b) UMAP dimensionality reductions of the scRNA-seq data on microglia and MG before and after microglia ablation. **(c)** Heatmaps of top predicted ligand scores from Lignature and three other methods, together with detection rate of the ligands in microglia and the maximum detection rate of their cognate receptors in MG before microglia ablation. **(d)** LR interaction scores were calculated by CellPhoneDB and NATMI based on the expression of ligand genes in microglia and the expression of the receptors in MG, and each ligand was scored by the maximum interaction score among the LR pairs between the ligand and its receptors. CellPhoneDB and NATMI-induced ligand scores, as well as gene-level Lignature predicted ligand scores zeroed out by detection rates of the ligands and their receptors at 0.3, were plotted in scatter plots against the detection rates of the ligands in microglia. Ligands with higher prediction scores than TNF and detection rates below 0.1 in microglia were labeled as false positives (FPs).

Application of Lignature to the scRNA data in MG predicted TNF was ranked as the top ligand (green dots in Figure 5c). In contrast, when we used NicheNet to assess the activity scores of the same ligands, TNF was positioned number 37. Several ligands highly ranked by NicheNet, such as IL1A, GIP, IL17F, C5, and LIF, were assigned lower rankings by Lignature. Upon examining the expression levels of these ligands in the sender cells (microglia) and their receptor’s expression in MG (before microglia ablation), we found that either the ligands or their receptors were undetectable or detected in a very limited portion of the cells, indicating they are unlikely to be responsible for the transcriptomic changes in MG. Moreover, the receptor differential expression-based method iTALK and a receptor expression-based method (max) ranked TNF number 148 and 42, failed in identifying TNF as an active ligand for the transcriptomic changes in MG.

We next used CellPhoneDB and NATMI, two gene expression-based methods for predicting ligand-receptor interactions, to identify active ligands responsible for the transcriptomic changes in MG. The ligands were ranked by the maximum interaction score between the ligands and their receptors. TNF was ranked number 17 by CellPhoneDB and number 7 by NATMI (Figure 5d). Interestingly, many ligands ranked higher than TNF by CellPhoneDB and NATMI, such as APOE, RPS19, YBX1, GRN, and GNAS were highly expressed in MG. Since we aimed to predict the ligands responsible for the transcriptomic changes in MG before and after microglia ablation, one would reasonably expect the active ligands to be mainly sourced from microglia. Hence the higher ranked ligands that were highly expressed in MG were less likely to be the ligands inducing the observed expression changes, compared to the experimentally validated ligand TNF, which was barely expressed in MG (detection rate < 0.002) but highly expressed in microglia. This analysis suggests that gene expression is not sufficient for the identification of active ligands, while in practice, incorporating the expression levels of both ligands and receptors, such as filtering the candidate ligands by the gene expression levels, would help reduce noises (Figure 5d).

### Usage of Lignature

We demonstrate the usage of Lignature by applying a scRNA-seq dataset of immune cell populations in checkpoint blockade-associated colitis and control samples^38^. We focused on the transcriptomic changes between samples from checkpoint inhibitor treated melanoma patients with and without histologically colitis, labeled “colitis” and “NC”, in myeloid, with all cell types in the dataset as potential ligand sending cells (Figure 6a, b).

**Fig.6.**
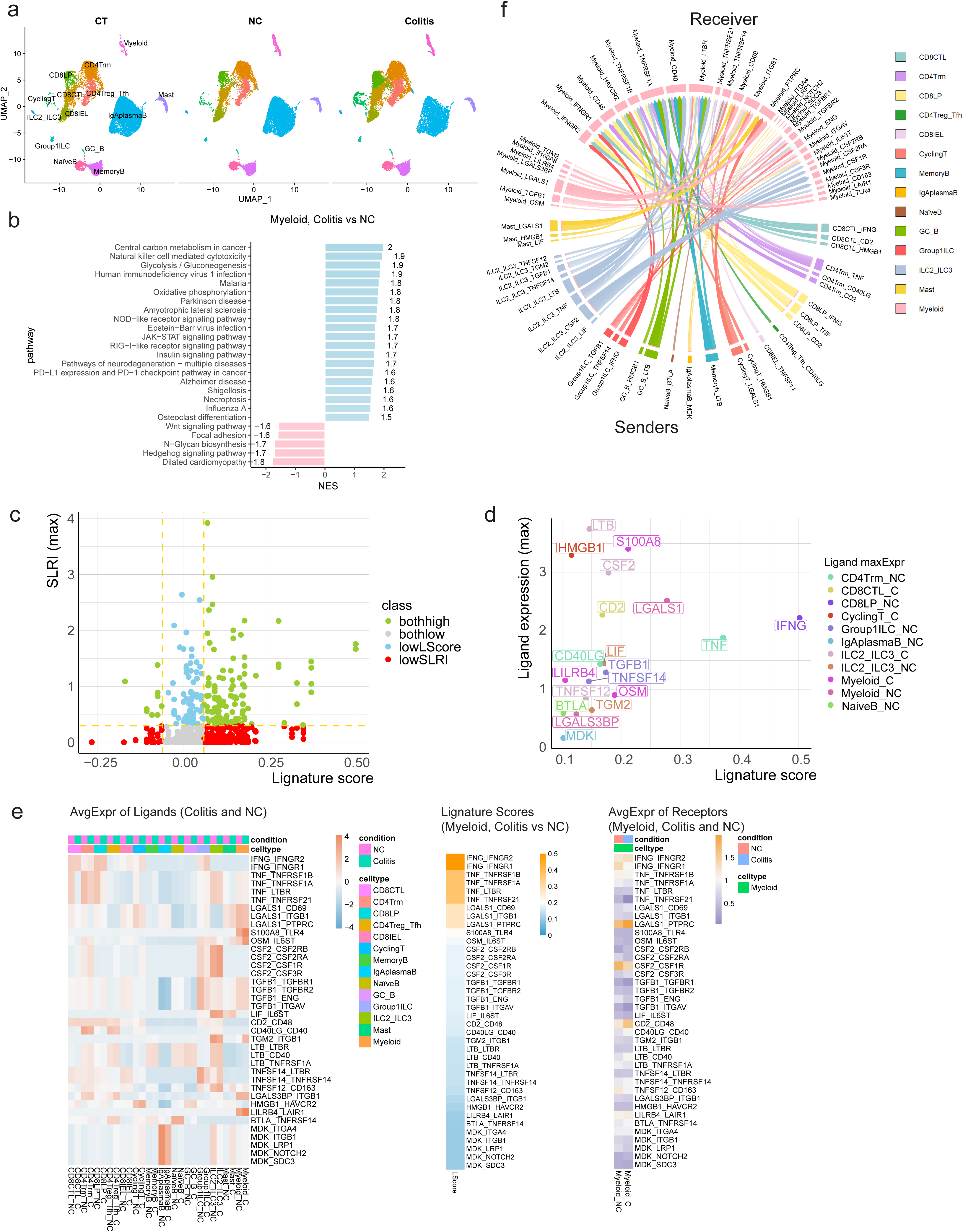
Application of Lignature on checkpoint blockade-associated colitis scRNA-seq dataset. **(a)** UMAP dimensionality reductions show cell types identified in the scRNA-seq data of immune cell populations in checkpoint blockade-associated colitis and control samples. **(b)** Enriched up- and down-regulated KEGG pathways in myeloid cells between colitis (checkpoint inhibitor treated melanoma patients with histologically colitis) and NC (checkpoint inhibitor treated melanoma patients who underwent endoscopic evaluation for suspected checkpoint inhibitor-induced colitis but were found to have a normal colonic mucosa endoscopically and histologically) samples. **(c)** Scatter plot of ligand similarity scores predicted by Lignature (x-axis) and the maximum LR interaction strength scores calculated by the method NATMI across all sender cells and conditions (y-axis) for all LR pairs with these scores available in the dataset. **(d)** Scatter plot of similarity scores predicted by Lignature (x-axis) and the maximum average expression levels across all sender cells and conditions (y-axis) of the ligands of identified LR interactions (detection rate of the ligand genes above 0.25 in any cell type, the detection of the receptor genes above 0.25 in myeloid and the ligand scores above 0.1). **(e)** Heatmaps of ligand scores predicted by Lignature (middle), the average expression of the ligands in all cell types identified in the colitis and NC samples (left), and the average expression of their receptors in myeloid cells in the colitis and NC samples (right), for LR interactions filtered by the criteria as in (d). **(f)** Circos plot of maximum LR interaction strength scores calculated by NATMI across the colitis and NC samples, of the LR pairs from all cell types to myeloid, with detection rates of both the ligand and receptor genes above 0.25, ligand scores above 0.1, and ligand expression above 75% of all cell types.

Our program can take the Seurat object, specified receiver and sender cell types, and a pair of sample conditions as input. Lignature scores the candidate ligands based on users’ choice of the similarity measurement metric (e.g., Pearson correlation coefficient, Spearman correlation coefficient, Cosine Similarity, Euclidean distance, etc.). In addition, we can also calculate the LR interaction score using specified scoring methods (e.g., CellPhoneDB, NATMI, or SingleCellSignalR). Lignature can help users select active ligands using both signature similarity scores and LR interaction strength scores (Figure 6c). Lignature provides multiple options for filtering the list of LR pairs (e.g. gene detection rate, average log2FC, adjusted p.value, ligand score, LR interaction strength score, LR interaction specificity, etc.) and produces various plots displaying enriched pathways, inferred ligands, and the associated LR interactions, providing intuitive visualizations for the analysis (Figure 6b-f and SupplFigure 4, before and after taking differential expression of the ligand and receptor genes into consideration). Through this analysis, Lignature successfully identified active ligand signaling to myeloid cells such as IFNG and TNF from T cells, as well as potential therapeutic targets for checkpoint inhibitor-induced colitis such as TNF.

## DISCUSSION

Understanding cell-cell communication is fundamental in decoding complex biological processes, and ligand-receptor interactions play a pivotal role in mediating these signals. In this study, we introduced Lignature, a comprehensive database of ligand signatures derived from experimental data, coupled with a predictive framework for identifying active ligands based on observed transcriptomic changes. By leveraging this resource, we address several limitations of existing computational methods, such as high false-positive rates from gene expression-based approaches and biases introduced by incomplete signaling network models.

Compared to network-based methods such as NicheNet, Lignature captures not only the target genes of the ligands, but also the direction (i.e., up- and down-regulation) and the magnitude of the expression changes of the target genes. Furthermore, it also captures cell type- and condition-specific transcriptomic changes. Our findings demonstrate that Lignature consistently outperforms existing methods, including NicheNet and iTALK, by providing more accurate predictions of active ligands. For example, in both controlled *in vitro* datasets and more complex *in vivo* scRNA-seq datasets, Lignature was able to rank biologically relevant ligands, such as TNF-α and SHH, at the top.

Despite these strengths, there are limitations that warrant further discussion. While Lignature incorporates a broad library of ligand signatures, its coverage is inherently dependent on the availability and diversity of experimental datasets. Certain ligands, particularly those with sparse experimental data, may be underrepresented, potentially limiting predictions in specific contexts. Furthermore, while the platform currently focuses on transcriptomic changes, integrating multi-omics datasets, such as proteomics or epigenomics, could provide a more comprehensive view of ligand signaling.

In summary, Lignature represents a significant advancement in the field of cell-cell communication, providing a robust, experimentally validated resource for inferring active ligand signaling. Its ability to address the limitations of existing computational methods while offering mechanistic insights underscores its potential as a valuable tool for both basic and translational research.

## METHODS

### Datasets collection

We collected 859 human ligands based on our previous work LRLoop^12^. We then developed an automatic pipeline to search Gene Expression Omnibus (GEO)^23^ for datasets with an experimental design that would potentially contain transcriptomic responses to each ligand.

Specifically, our collection of the relevant datasets includes the following three steps. We started by the list of all 25,868 platform accession identifiers downloaded from GEO and collected the metadata for each of them using the R package “GEOquery”^39^, filtered the list by restricting the organism to “Homo sapiens”, and collected the list of 97,693 data-series identifiers corresponding to the 6,278 filtered platforms, where each data-series could include multiple datasets. Then again using GEOquery, we curated metadata for the collected data-series including data “type”, “organism”, “molecule”, “platform id”, “cell line”, “cell type”, “tissue”, as well as data “title”, “abstract”, “url”, “geo accession”, “overall design”, “sample taxid”, “summary”, etc. The list of data-series was then filtered by requiring the “molecule” entry of the corresponding metadata to include the strings “total RNA”, “polyA RNA” or “nuclear RNA”, resulted in a list of 72,738 data-series identifiers for further systematic searching of ligand and experiment keywords.

In order to search the curated metadata of the GEO series systematically for potential ligand signatures, we collected a set of “ligand keywords” including gene full names, symbols, and aliases for each ligand, as well as a set of 105 experiment keywords such as “ligand”, “receptor”, “treatment”, “stimulate”, “activate”, “expose”, “block”, “overexpress”, “knockout”, “knockdown”, “silence”, “inhibit” and “in the presence of”. The curated metadata of the GEO series list was then queried with the ligand keywords and experiment keywords. This automatic searching process reduced the number of candidate data-series in the list to 9,187 for manual curation.

Finally, after careful examination of the description of each of the collected data series in GEO, we removed the ones that did not serve our purpose and obtained a list of 732 data-series, containing transcriptomic response data for 281 ligands that were not included in the database CytoSig^21^ (Figure 1a).

In addition, to create a comprehensive ligand signaling signature database, we took advantage of CytoSig^21^ and ImmuneDictionary^22^ by including the cytokine signatures from these sources into Lignature, and included the signatures of the 90 test bulk datasets after method assessment by these test datasets.

### Data processing

We have two major data types: gene expression array and RNA-seq datasets. For Affymetrix array data with raw data .cel files provided, we adopted the “maEndtoEnd”^40^ workflow, which generates and preprocesses expression data using the R package “Oligo”^41^ and perform differential expression analysis using the “limma”^42^ package. For other forms of array data, we used processed data and performed differential expression analysis by “limma”. Briefly, after RMA calibration using the “oligo::rma” function, an intensity-based filtering was performed by removing transcripts without intensities higher than a threshold of two in at least the number of arrays of the smallest experimental group, followed by annotation of the transcript clusters with gene symbols and removal of multiple mappings. Subsequently, a contrast matrix was constructed using functions “model.matrix” and “makeContrasts” and used as input of a standard differential expression analysis pipeline in limma, i.e., “lmFit”, “contrasts.fit”, and “eBayes”. Then for each gene, the most significant ProbeSet corresponding to it with the lowest adjusted p.value was identified and used as the representative of that gene.

RNA-seq datasets were processed with a combination of “edgeR”^43^ and “limma”. In brief, a DGEList object was created using the function “DGEList” with the expression count matrix, genes were annotated by gene symbols. and lowly expressed genes were identified for removal by the “filterByExpr” function with default settings. Following gene expression normalization through “calcNormFactors” with the “TMM” method, the standard limma-voom pipeline was applied for differential expression analysis.

After the differential expression analysis, different types of gene identifiers were converted to gene symbols, and the vectors of log2FC values of expressed genes were collected as the gene-level ligand signaling signature. For datasets with multiple treatment conditions, e.g., durations or doses, a separate signature was created for each different treatment condition.

In addition to the gene-level response signatures, we also calculated KEGG pathway signatures for each ligand. Specifically, we collected gene sets of 302 KEGG pathways with 10 ∼ 500 genes by the R package “graphite”^44^. Then for each gene-level signature, we pre-sorted the log2FC vector in descending order and performed Gene Set Enrichment Analysis using the R package “fgsea”^45^, calculated the Normalized Enrichment Score (NES) for the KEGG pathways. The vectors of NES were collected as the pathway-level signature (Figure 1a).

### Prediction of acting ligands based on ligand signatures

Using the Lignature database as a reference for ligand signaling responses, we provided a companion computational tool that infers the ligands responsible for transcriptomic changes within receiver cells and the corresponding cell-cell interaction networks between multiple cell types or clusters. For a dataset of interest, taking matrices resulted from differential expression analysis, or the Seurat^46^ object of single cell RNA-seq datasets as input, Lignature scores each ligand signature in the database by the similarity compared to the signature of the input data. Options of the similarity measurement metrics include Pearson correlation coefficient, Spearman correlation coefficient, cosine similarity, Euclidean distance, Manhattan distance, and the Euclidean or Manhattan distance normalized by vector lengths. Users can filter or zero out the input data and the reference ligand signatures by detection rates, log2FC values or adjusted p.values. Subsequently, each ligand is scored by its corresponding maximum signature score or the signature score with the highest absolute value in the database, based on user’s choice. Furthermore, Lignature provides multiple options for calculating expression-based LR interaction strength scores such as CellPhoneDB, NATMI, and SingleCellSignalR. In addition, users are provided the choices of filtering the LR-interactions by the ligand scores, expression or differential expression of the ligands and their receptors, interaction strength scores and the interaction-specificities. Lastly, Lignature features the options to visualize predicted ligand scores, expression of the ligands and their receptors, as well as inferred strengths of LR interactions, providing an intuitive summary of the predicted LR activities across different conditions (Figure 1c).

### Comparison with other methods

Five other different methods of ligand activity prediction were compared to Lignature. Specifically, for each test dataset, “NicheNet” ligand activity scores were calculated by the R package “nichenetr” according to its recommended parameter settings, i.e., expressed genes with |log2FC| >= 1 and adjusted p.value < 0.1 were used as differentially expressed genes; The method “iTALK [maxRlfc]” scored each ligand by the maximum absolute log2FC value among the cognate receptors; “iTALK [meanRlfc]” scored each ligand by the arithmetic mean of its receptors’ absolute log2FC values; “Receptor expression [max]” scored each ligand by the maximum expression value of the cognate receptors; and “Receptor expression [mean]” scored each ligand by the mean expression value of the cognate receptors. On the other hand, with Lignature, “Lignature [gene log2FC]” scored each ligand by Lignature with the gene-level signatures, while “Lignature [pathway NES]” scored each ligand by Lignature using the pathway-level signatures, with the vector similarities measured by the Pearson correlation coefficients.

Known treatment ligands were labeled “TRUE” and other candidate ligands were labeled “FALSE”. Ranks of the true treatment ligands were plotted in Figure 3c. In Figure 3d, we present the AUC values calculated from ROC curves of the methods under comparison for each test dataset separately. ROC curves of methods under comparison from test results of all 90 test datasets combined are presented in Figure 3e.

This assessment was performed using our initial collection of signatures, and the 90 test datasets were organized and added to Lignature afterwards.

## DATA AND CODE AVAILABILITY

All data analyzed in this study are included in the published article, supplementary files, and publicly available repositories. Lignature database is available at https://github.com/yingxinac/LignatureData, and the companion R package is at https://github.com/yingxinac/Lignature.

## ACKNOWLEDGEMENT

This study was supported by National Eye Institute (NEI) grant R01EY031779 (JQ).

## DECLARATION OF INTERESTS

The authors declare no competing interests.

## Figure legends

**Suppl.Fig.1.**
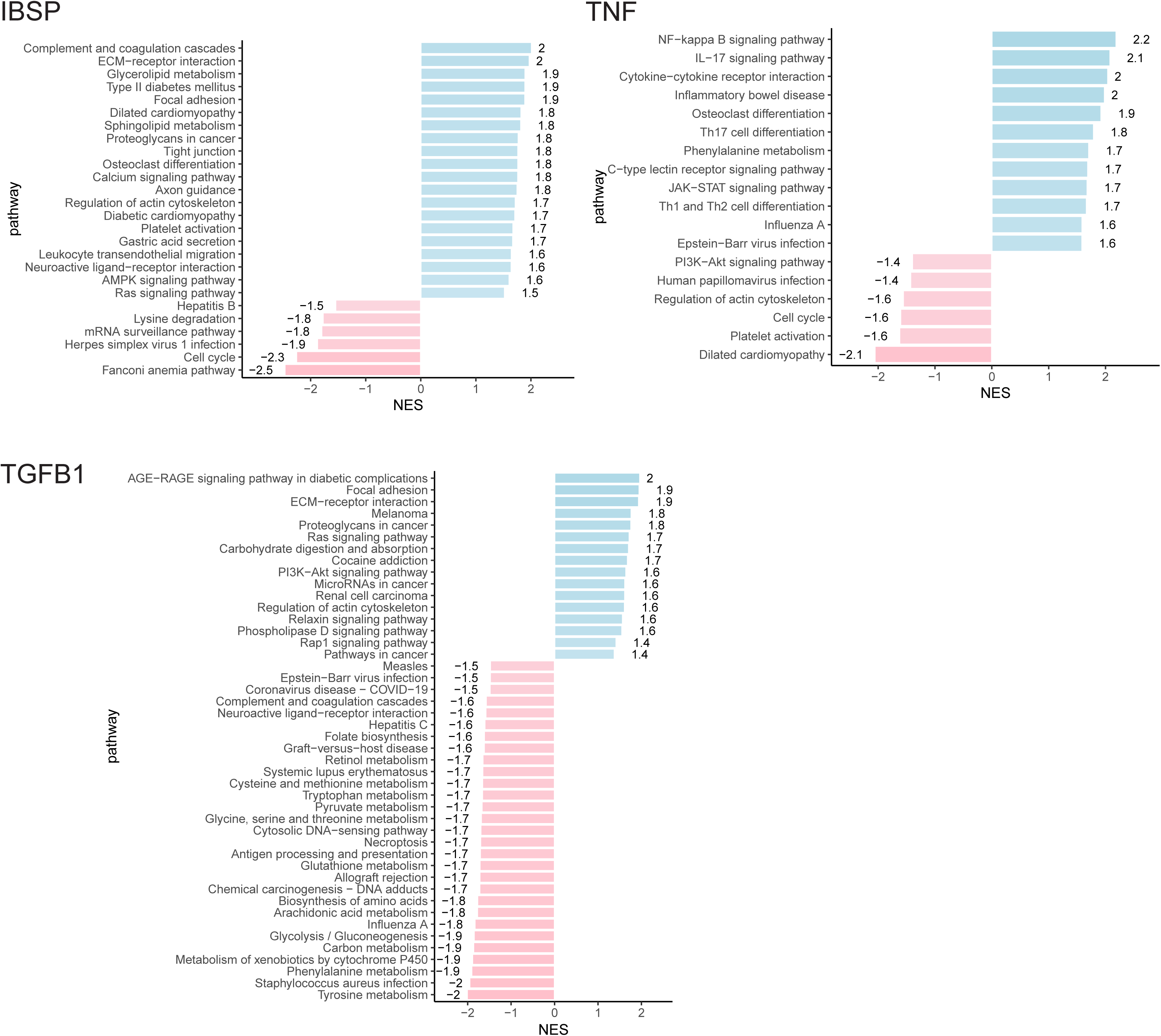
Additional examples of pathway enrichment in ligand signatures. Examples of enriched up- and down-regulated KEGG pathways in sample ligand signatures of IBSP, TNF, and TGFB1.

**Suppl.Fig.2.**
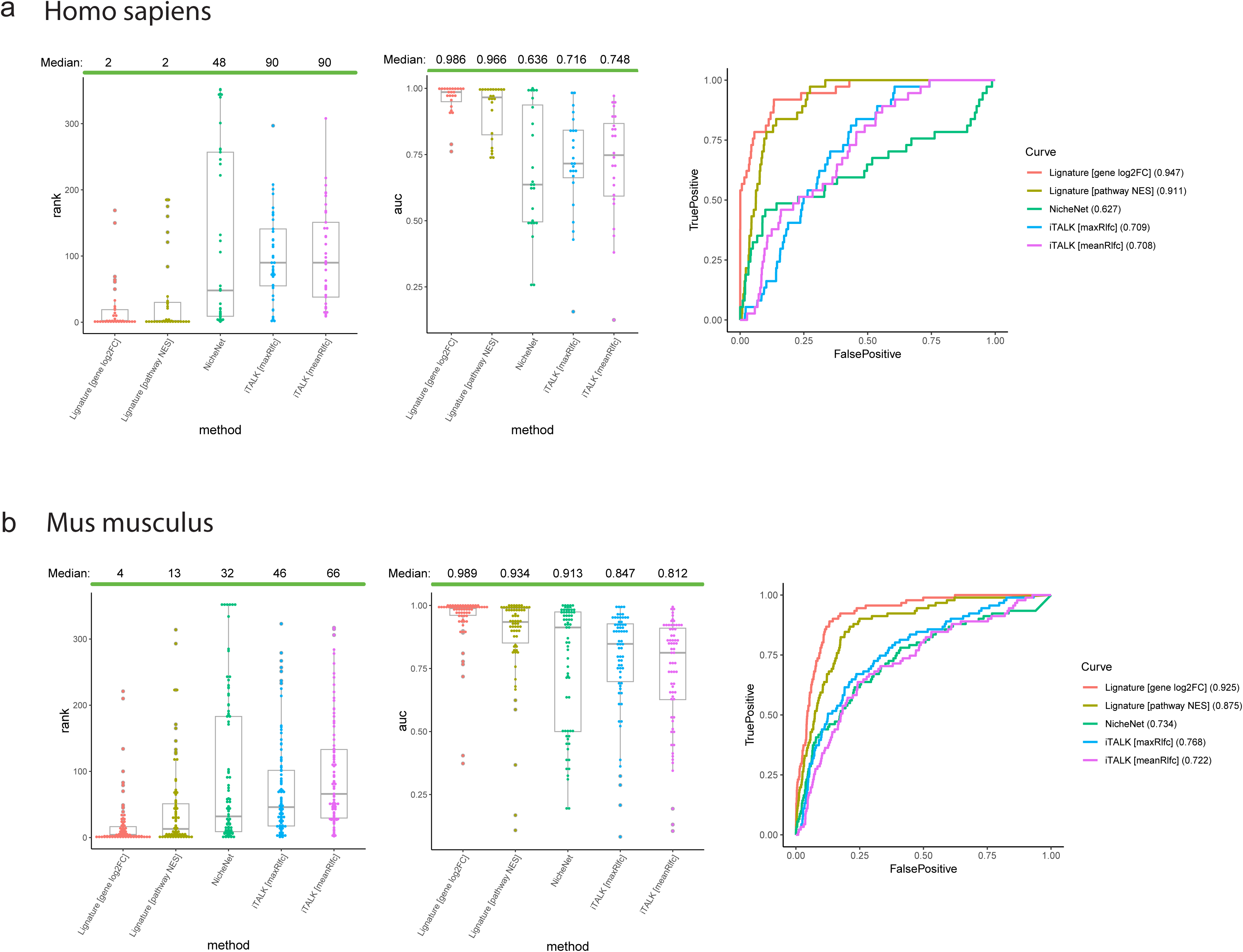
Comparison of Lignature to other methods in signaling inference **with (**a**) human and (**b**) mouse bulk in vitro datasets separately.**

**Suppl.Fig.3.**
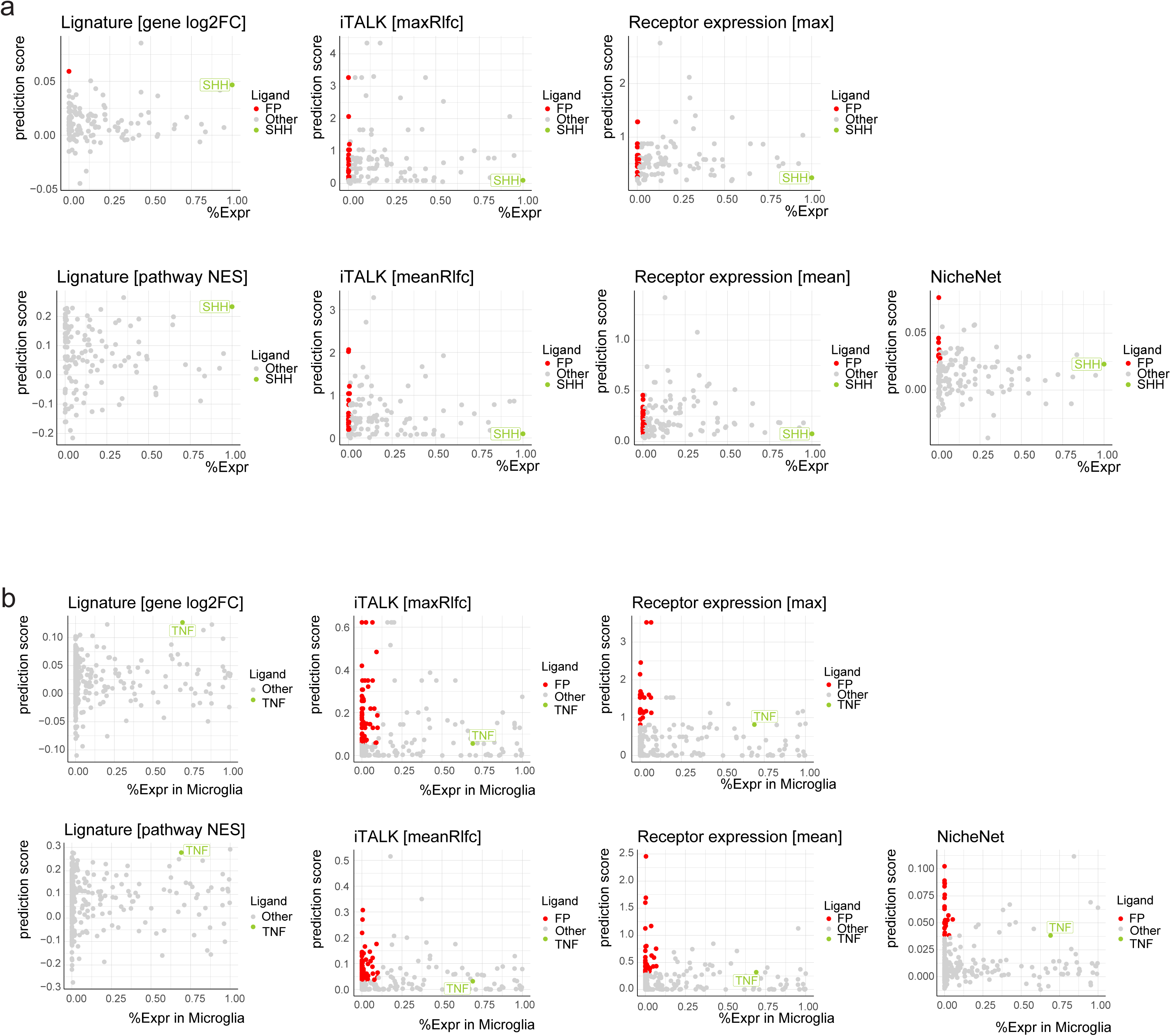
(a) Scatter plots of predicted ligand scores and the maximum detection rates for SHH-treated and untreated samples across all cells of candidate ligands for each method. Ligands with higher prediction scores than SHH but a detection rate below 0.01 across all cells were labeled as false positives (FPs). SHH was assigned a detection rate of 1 since recombinant SHH was externally added to the culture without clear sender cells. **(b)** Scatter plots of predicted ligand scores against detection rates in microglia of candidate ligands for gene-level and pathway-level Lignature, together with five other methods. Ligands with higher prediction scores than TNF and a detection rate below 0.1 in microglia were labeled as false positives (FPs).

**Suppl.Fig.4.**
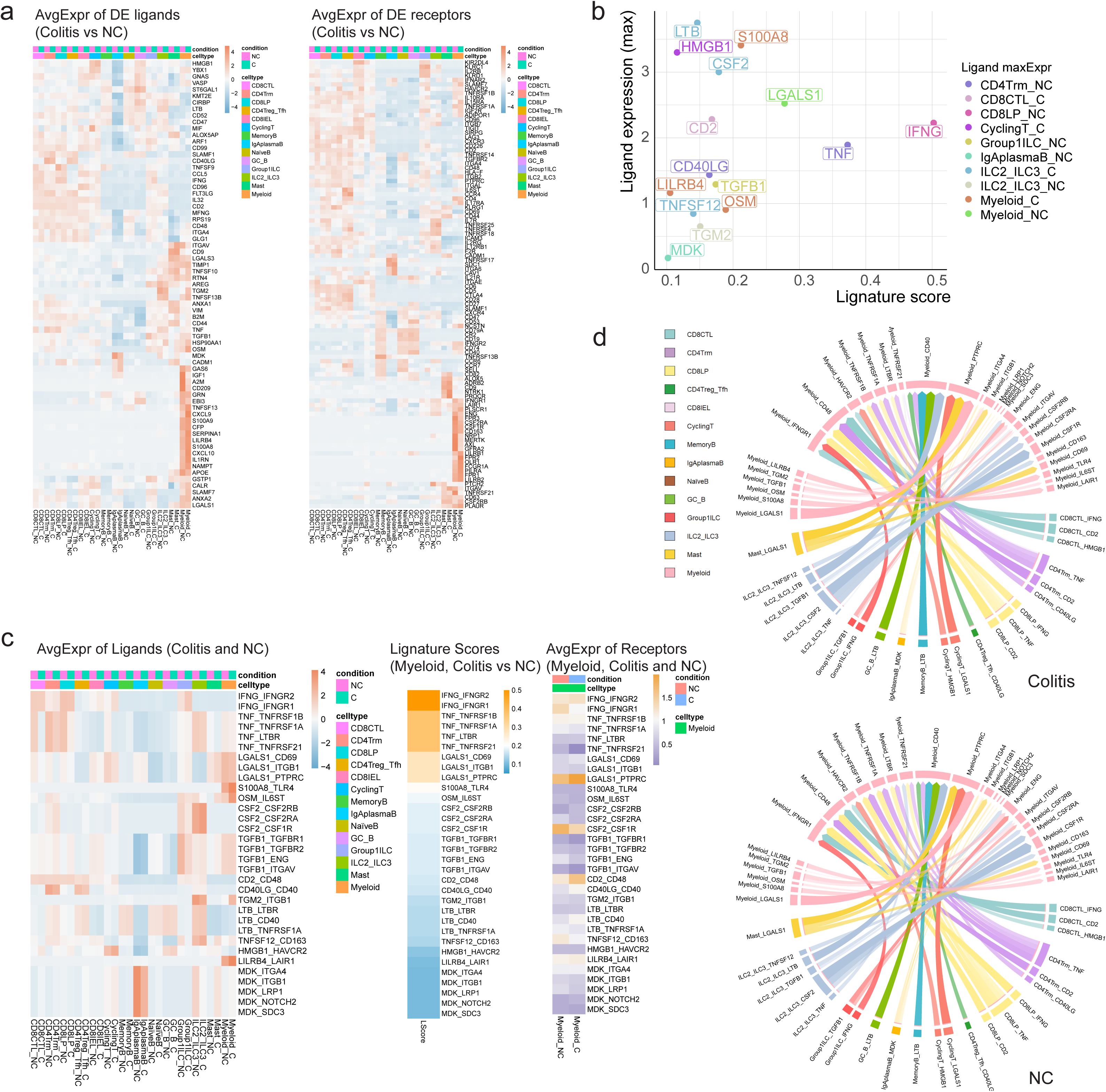
Lignature identifies active differential LR interactions to myeloid in checkpoint blockade-associated colitis. **(a)** Heatmaps of the average expression of differentially expressed ligand genes and receptor genes (average log2FC above 0.2 and adjusted p.value below 0.01) in the colitis and NC samples. **(b-d)** Analysis results of Lignature from further filtering the LR interactions by differential expression of the ligand or receptor genes.

## Notes

### Competing Interest Statement

The authors have declared no competing interest.

